# Pythia 2.0: New Data, New Prediction Model, New Features

**DOI:** 10.1101/2025.03.25.645182

**Authors:** Julia Haag, Alexandros Stamatakis

**Affiliations:** Computational Molecular Evolution Group, Heidelberg Institute for Theoretical Studies, Heidelberg, Germany; Biodiversity Computing Group, Institute of Computer Science, Foundation for Research and Technology - Hellas, Heraklion, Crete, Greece; Institute for Theoretical Informatics, Karlsruhe Institute of Technology, Karlsruhe, Germany

## Abstract

Maximum Likelihood (ML) based phylogenetic inference is time- and resource-intensive, especially when initiating multiple independent inferences from distinct comprehensive tree topologies. Performing multiple independent inferences is often required to (sufficiently) explore the vast search space of possible unrooted binary tree topologies. Yet, these independent inferences do not necessarily converge to a single phylogeny or at least topologically highly similar trees. While for *easy-to-analyze* multiple sequence alignments (MSAs), one is likely to obtain a conclusive, single phylogeny, *difficult-to-analyze* MSAs yield topologically highly distinct, yet statistically indistinguishable tree topologies. In 2022, we proposed a compute-intensive approach to quantify the inherent *difficulty* of a phylogenetic analysis for a specific, given MSA, and also trained a machine-learning based prediction model called *Pythia* to substantially reduce the computational cost of determining the difficulty. Pythia can predict the difficulty for a given MSA with high accuracy, while being substantially faster than even a single ML tree inference. Pythia predicts the difficulty on a scale from 0 (easy) to 1 (difficult). Here, we present all improvements to Pythia that we have introduced since our initial publication in 2022. We trained a new prediction model using approximately three times more MSAs and a new type of machine learning model. We improved the runtime of two feature computations, and we also introduced two additional prediction features. Our latest version Pythia 2.0 is slightly more accurate than our initial version and is also approximately twice as fast. Finally, we also present and make available, the novel and easy-to-use command line tool *PyDLG* that allows to compute the ground-truth difficulty seamlessly for a given MSA. This ground-truth difficulty can be used, for instance, as a prediction target for training a new Pythia model.

## 1 Background

In 2022, we published our work on predicting the *difficulty* of phylogenetic analyses of multiple sequence alignments (MSAs). We introduced a difficulty quantification that captures the potential inherent “ruggedness” of the Maximum Likelihood (ML) surface on a scale from 0 (*easy*) to 1 (*difficult*). An *easy* MSA exhibits a clear likelihood peak, and multiple independent ML tree inferences will converge to highly similar tree topologies. In contrast, multiple independent tree inferences on a *difficult* MSA will yield highly distinct trees that are statistically indistinguishable from each other by standard likelihood-based statistical tests for phylogenetics. In addition to quantifying this phenomenon, we also published *Pythia*, a Random Forest regression model that can accurately predict the difficulty of a given MSA, based on eight fast-to-compute prediction features. We showed that Pythia is substantially faster than inferring even a single ML tree. We therefore suggested that researchers predict the difficulty of their MSA using Pythia prior to setting up and conducting any phylogenetic analyses. This does not only allow for an informed decision on the required analysis setup (for instance, how many trees to infer and what type of post-analyses to conduct), but might also prevent potentially time- and resource-intense analyses that are unlikely to yield a conclusive phylogeny. Building on Pythia, and a thorough performance comparisons of various ML tree inference heuristics by Hoehler *et al*. [6], Togkousidis *et al*. [16] implemented an adaptive ML tree inference procedure for the popular RAxML-NG (*adaptive RAxML-NG*) ML tree inference tool. Adaptive RAxML-NG predicts the difficulty of an MSA using Pythia and subsequently categorizes the MSA into *easy* (difficulty ranging from 0.0 to 0.3), *intermediate* (0.3 to 0.7), and *difficult* (*>* 0.7). Depending on the difficulty category, adaptive RAxML-NG adjusts the default number of inferred trees, and also executes an appropriately adapted tree search strategy. Compared to *standard* RAxML-NG by Kozlov *et al*. [10], adaptive RAxML-NG exhibits an average speedup of 16× on easy as well as difficult MSAs, and of 1.8× on intermediate MSAs. Note that the speedup cannot exclusively be attributed to the difficulty-induced changes, as Togkousidis *et al*. [16] also adjusted the difficulty-independent tree inference algorithm to allow for faster tree inferences. Yet, adaptive RAxML-NG demonstrates the substantial potential of integrating Pythia-informed strategies into phylogenetic analyses pipelines.

Here, we present the changes we introduced in Pythia over the past three years. We included substantially more MSAs in our training dataset and introduced support for morphological data in addition to DNA and amino acid (AA) data. In more recent versions of Pythia, we replaced the Random Forest regression model with a *Gradient Boosted Trees* regressor, yielding improved prediction performance. We also included two additional prediction features, and were able to reduce the number of inferred maximum parsimony (MP) trees (the main computational bottleneck of feature computation) required for obtaining two of the already existing features. Overall, these changes induce a slight prediction accuracy improvement. The mean absolute error (MAE) improved from 0.09 to 0.08, and the mean absolute percentage error (MAPE) from 2.9% to 2.2%. Additionally, our most recent Pythia version is more than twice as fast compared to the initial published version. In the following, we provide detailed explanations of all changes, as well as respective performance analyses and comparisons. Finally, we also present a novel and easy-to-use command-line tool that allows users to compute the “true” difficulty of an MSA based on our quantification presented in Haag *et al*. [5]. This might serve for training Pythia on novel data types, such as data from linguistics or potential alignments of protein structure alphabets.

## 2 Pythia 2.0

In the following, we refer to our first version of `Pythia` as `Pythia 0.0`. This denotes the version of our difficulty predictor at the time of the initial publication. We refer to the latest version at the time of this publication as `Pythia 2.0`. This version corresponds to the Pythia version available as release v2.0.0 on GitHub (https://github.com/tschuelia/PyPythia/releases/tag/2.0.0).

### 2.1 New Data

We trained `Pythia0.0` on 3250 empirical MSAs obtained from TreeBASE [14] comprising 74% DNA MSAs and 26% amino acid (AA) MSAs. For our latest predictor `Pythia 2.0`, we increased the training dataset to 10 461 MSAs, using MSAs from TreeBASE and the RAxML Grove database [7]. Note that the publicly available RAxML Grove database contains phylogenetic trees only. The underlying, fully anonymized MSAs are only available internally within our research group, and we only use these MSAs to compute the features required for training Pythia. This new training dataset comprises 87% DNA, 9% AA, and 4% morphological data. Since we explicitly include morphological MSAs, `Pythia 2.0` can now accurately predict the difficulty for this data type as well. Note that we provide a single predictor for all data types, as we observe no substantial differences in prediction accuracy when using a combined predictor compared to using an independent, dedicated predictor for each data type separately (see below).

### 2.2 New Prediction Model

The prediction model we used in `Pythia 0.0` was a Random Forest regression model [2]. Since then, we transitioned to using a Gradient Boosted Tree (GBT) regressor [4] instead, as we observe an improved performance (see below). In analogy to a Random Forest, a GBT comprises an ensemble of Decision Trees. While the Decision Trees in a Random Forest can be trained independently, and thus in parallel, the Decision Trees in a GBT are trained sequentially. Each individual Decision Tree strives to improve upon the overall prediction by attempting to predict the prediction error of the preceding Decision Tree. Consequently, training a GBT regressor is, in general, more time-consuming than training a Random Forest regressor. Yet, the prediction using a GBT can be parallelized since the individual Decision Trees can be queried independently after training. Thus, the prediction time of GBT and a Random Forest regressors are comparable (given the same number of Decision Trees, each with the same maximum tree depth).

In `Pythia 2.0`, we rely on the implementation of GBTs in the *LightGBM* Python package [8]. We optimized the hyperparameters of the GBT regressor using the *Optuna* framework [1].

To ensure a fair and unbiased performance evaluation, it is important to only evaluate the trained model on data that has not been used for training. With `Pythia 0.0` we used an 80-20 train-test split approach for evaluation, that is, we used 80% of the MSAs for training, and the remaining 20% for evaluation. With `Pythia 2.0`, we evaluated the performance using a 10-fold cross validation approach. This means that we first split the training dataset into 10 equally sized, disjoint MSA subsets. We subsequently trained a regression model using 9 out of 10 subsets, and evaluated the trained model on the remaining subset. We repeated this procedure 10 times, such that each of the 10 subsets was used for model evaluation exactly once. Finally, we obtained the overall prediction performance of `Pythia 2.0` as average performance over all 10 folds. We evaluate and compare `Pythia 0.0` and `Pythia 2.0` using the Mean Absolute Error (MAE) and Mean Absolute Percentage Error (MAPE). We compute the MAE and MAPE as

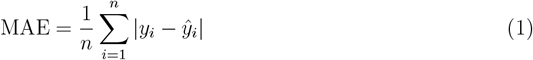

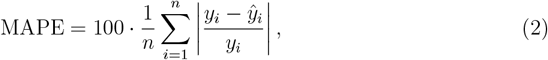

where *n* is the number of MSAs used for training, *y*_*i*_ is the ground-truth difficulty of the *i*-th MSA, and 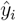 the respective predicted difficulty. The best possible value for both evaluation metrics is 0.

### 2.3 New Features

`Pythia 0.0` relied on 8 prediction features for each MSA. Four features capture intrinsic attributes of the MSA: *sites-over-taxa* ratio, the *patterns-over-taxa* ratio, the proportion of invariant sites (% *invariant*), and the proportion of gaps (% *gaps*). Two features capture the information content of the MSA: the *Entropy* and the *Bollback Multinomial*. Finally, two features approximate the ruggedness of the ML tree space based on 100 trees inferred under the maximum parsimony (MP) criterion: the average pairwise RF-Distance between the inferred MP trees (*RF*_MP_) and the proportion of unique MP tree topologies 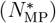. See Haag *et al*. [5] for further details of these features.

In `Pythia 2.0`, we include two additional features: the *patterns-over-taxa* ratio as an additional MSA attribute and the *Pattern Entropy* as an additional approach to capturing the information content of the MSA. The patterns-over-taxa ratio is simply the number of unique sites patterns in the MSA divided by the number of sequences (taxa). The Pattern Entropy is an entropy-like measurement based on the number and frequency of unique site patterns in the MSA. We compute the Pattern Entropy *H*_*P*_ as

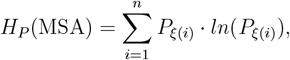

where ξ(*i*) denotes the *i*-th unique pattern and *P*_*ξ*(*i*)_ the number of times this pattern occurs. Note that the computation of this feature is related to the *Bollback Multinomial* feature. However, we expect it to be less biased by the total number of sites in the MSA.

#### 2.3.1 Reducing the Number of Maximum Parsimony Trees

To improve upon the runtime of Pythia’s feature computations, we aimed to reduce the number of MP trees to compute the *RF*_MP_ and 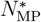 features. In `Pythia 2.0`, we rely on only 24 inferred trees, compared to the 100 trees used in `Pythia 0.0`. In the following, we detail the analyses leading to the conclusion that 24 constitutes the optimal number of MP tree inferences.

The MP criterion is 𝒩𝒫-hard [3]. Consequently, implementing tree inferences under this criterion requires heuristic algorithms. For instance, RAxML-NG implements a randomized stepwise addition order algorithm to infer MP trees. A consequence of these heuristics is that different initializations may result in topologically distinct inferred trees, and can consequently yield variations in the average relative pairwise RF-Distance between a set of inferred trees (*RF*_MP_). In `Pythia 0.0`, we inferred 100 MP trees, simply because we based the ground-truth difficulty on 100 ML trees. However, we observed that the runtime of `Pythia 0.0` is dominated by the runtime of the MP tree inference (approx. 50% of total runtime). Thus, to reduce the computational overhead of the MP-based features in `Pythia 2.0`, we aimed to reduce the number of inferred MP trees.

Our goal was to find the minimum number of MP trees *n*, such that the difference in *RF*_MP_ using only *n* trees compared to the *RF*_MP_ using 100 trees is lower than the variance in *RF*_MP_ using 100 trees inferred under varying random seeds.

To determine *n*, we performed the following analyses on 1000 MSAs that were randomly selected from our training dataset.

For each MSA, we first inferred 20 sets of 100 MP trees each, using a different random seed for each set. Then, for each seed *i*, we compute the RF-Distance between subsets of *m* = 3 … 99 trees sampled from the 100 trees inferred under seed *i*. We then consider two main metrics: the standard deviation of the absolute difference of RF distances between *m* trees and 100 trees across all seeds 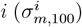, and the standard deviation of the absolute difference of RF distances among different seeds *i, j* for the same number of trees 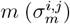. Finally, we compute the mean of these standard deviations across all MSAs and seeds for each *m*:

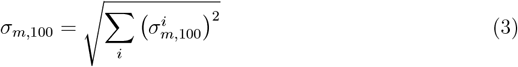

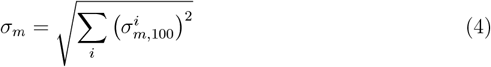

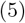

Note that the mean of a standard deviation is computed as the square root of the average variance.

Finally, we visualize both metrics as a function of the number of trees *m* in Figure 1. The x-axis shows the number of trees (*m* = 3 … 99), and the y-axis shows the average standard deviation for *σ*_*m*,100_ (solid, lighter trace) and *σ*_*m*_ (dotted, darker trace). The dashed horizontal line highlights *σ*_100_, corresponding to the variance between seeds when inferring 100 MP trees. We use this as a benchmark, since it represents the variance in *RF*_MP_ under different random seeds in `Pythia 0.0` which relies on 100 MP trees. Therefore, the intersection of *σ*_*m*_ with *σ*_100_ corresponds to the minimum number of MP trees such that the difference in *RF*_MP_ using only *m* trees compared to 100 trees is less than or equal to the variance of differences in *RF*_MP_ when using 100 trees but varying the random seed. As indicated by the dashed vertical line, for *m* = 23 *σ*_*m*_ < *σ*_100_. This means that inferring 23 MP trees constitutes the optimal tradeoff between approximation accuracy and runtime. We decided to instead use one additional tree and infer 24 MP trees, as 24 inferences are easily parallelizable in machines with different core counts (divisible by 2, 3, 4, 6, 8, and 12).

**Figure 1:**
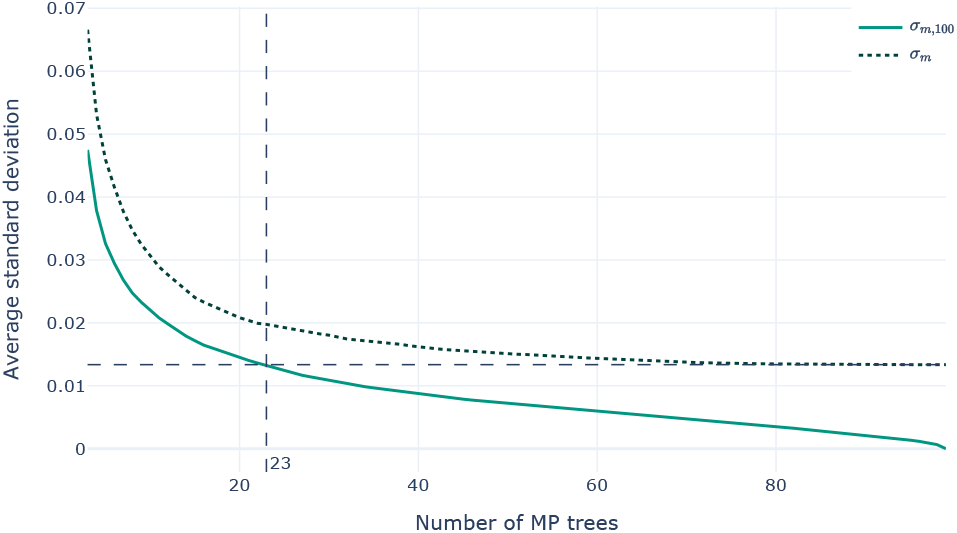
Visualization of *σ*_*m*,100_ and *σ*_*m*_ for *m* = 3 … 99 averaged across 1000 MSAs. The dashed horizontal line indicates *σ*_100_ and the dashed vertical line the intersection of *σ*_*m*_ with *σ*_100_ at 23.

## 3 Evaluation

### 3.1 Training Data Comparison

Figure 2 visualizes the distribution of ground-truth difficulty labels and feature values in the training datasets for `Pythia 0.0` and `Pythia 2.0`. For better visualization, we removed values above the 95th percentile for the patterns-over-taxa, sites-over-taxa, Entropy, and Pattern Entropy features. We removed values below the 5th percentile for the Bollback Multinomial feature.

**Figure 2:**
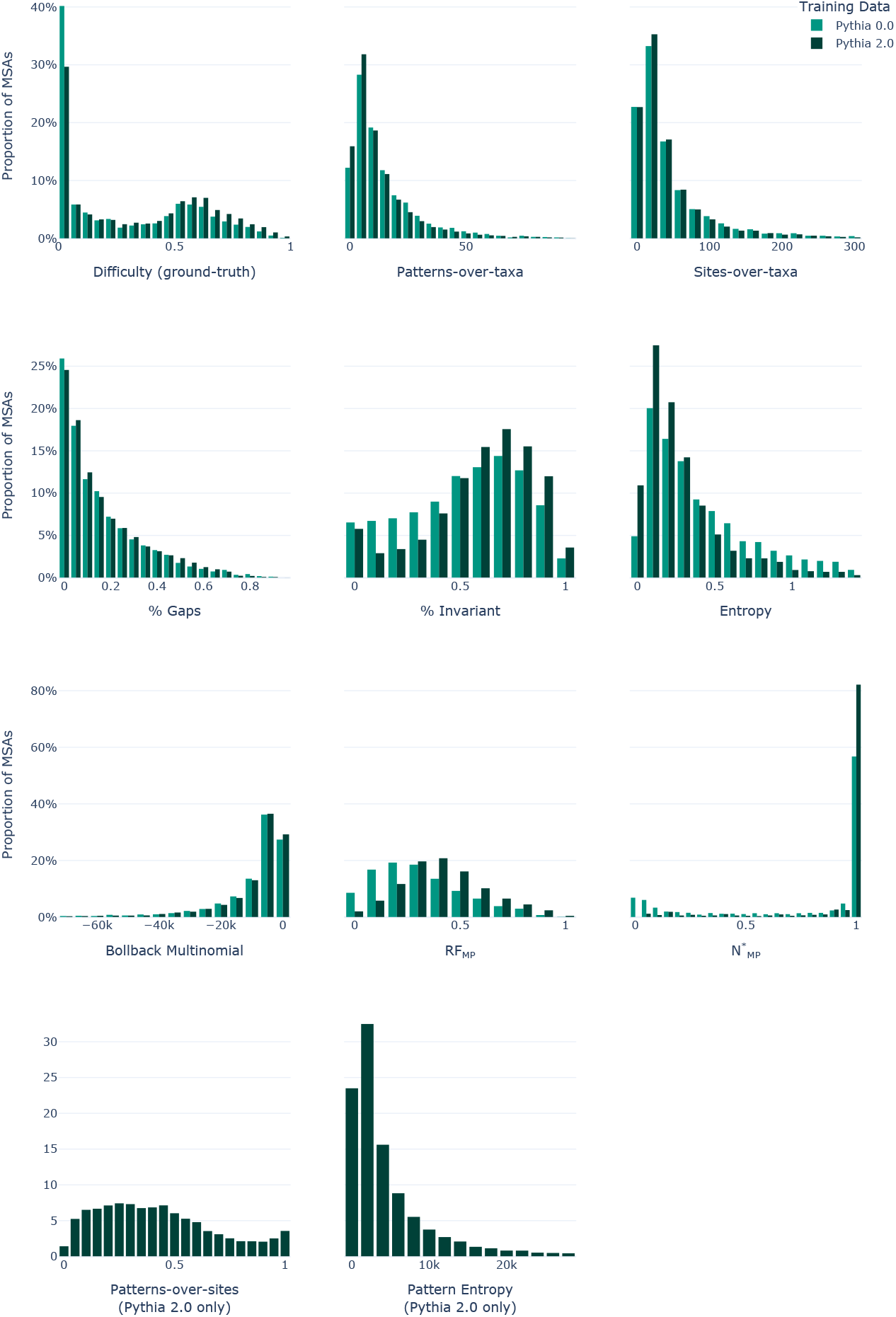
Distribution of ground-truth difficulty label and feature values in the training dataset for `Pythia 0.0` and `Pythia 2.0`. Note that we introduced the patterns-over-site and Pattern Entropy features in `Pythia 2.0`.

In the training data of `Pythia 0.0`, 46% of the MSAs are very easy, with a ground-truth difficulty of less than or equal to 0.1. Only 9% of MSAs are difficult (difficulty 0.7 or higher). With our training data for `Pythia 2.0`, we still observe an abundance of very easy MSAs (36%). We were nonetheless able to increase the number of difficult MSAs to 1401 (13%).

### 3.2 Model Comparison

#### 3.2.1 Prediction Performance

Pythia predicts the difficulty of an MSA on a scale of 0.0 (easy) to 1.0 (difficult). Our initial model `Pythia 0.0` has a MAE of 0.09 and a MAPE of 2.9%. With `Pythia 2.0` we slightly improved the prediction accuracy to a MAE of 0.08 and a MAPE of 2.2%. When analyzing the prediction error, we noticed that Pythia tends to overestimate the difficulty of MSAs with a difficulty ≤0.3 and underestimate the difficulty for MSAs with a difficulty *>* 0.3 (see Figure 3). We observe this effect with both `Pythia 0.0` and `Pythia 2.0`. However, with `Pythia 2.0`, this effect is less pronounced. We suspect that this constitutes an artifact of the uneven difficulty distribution in the training data. Thus, future work should focus on obtaining more intermediate and difficult MSAs.

**Figure 3:**
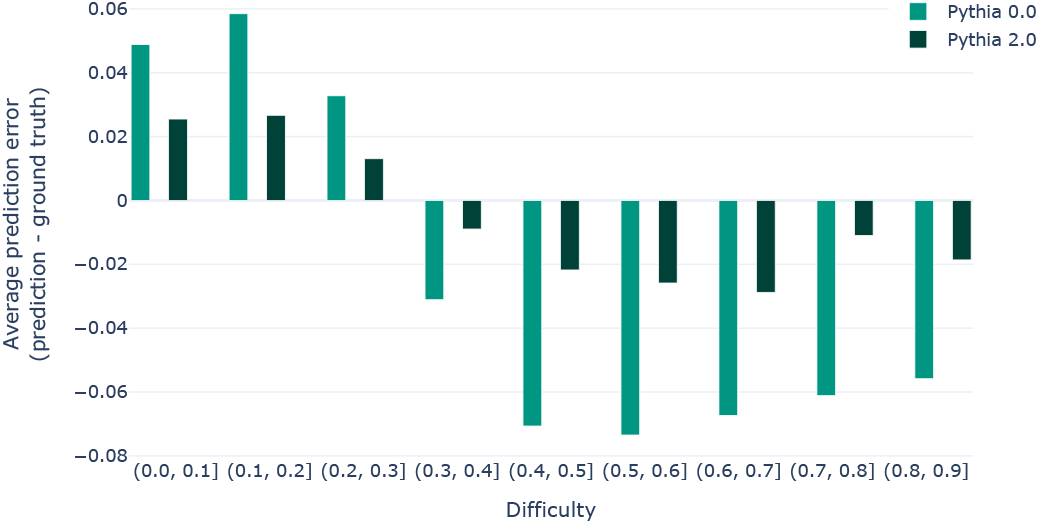
Average prediction error per difficulty range. The figure shows the error for `Pythia 0.0` and `Pythia 2.0` on their respective training datasets. We compute the prediction error as *predicted difficulty - ground-truth difficulty*.

We decided to train a single predictor for DNA, AA, and morphological data rather than training a separate predictor for each data type. To justify this design choice, we compare the prediction error of `Pythia 2.0` per data type separately. As Figure 4 shows, the overall performance of Pythia is comparable over all three data types.

**Figure 4:**
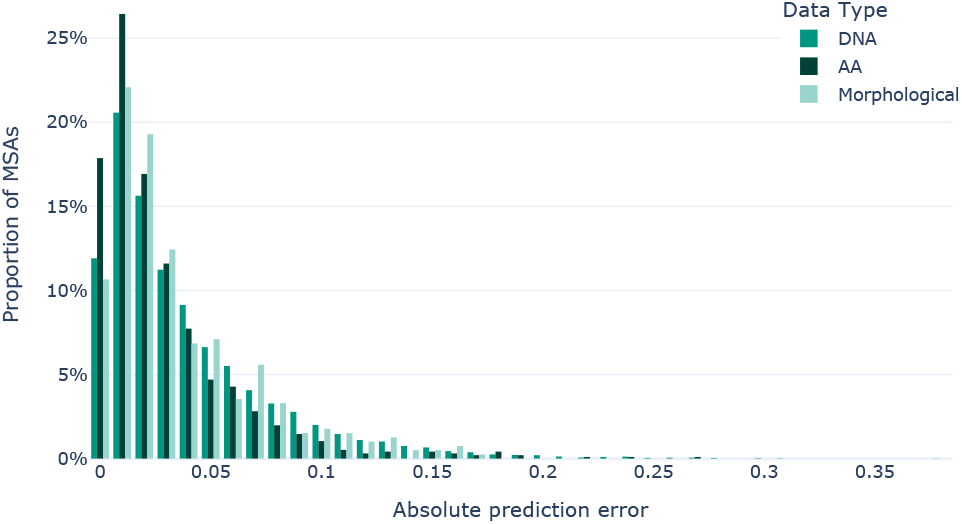
Average absolute prediction error of `Pythia 2.0` per data type.

To further justify this design choice, we also trained three separate predictors per data type (*f*_DNA_, *f*_AA_, *f*_M_), and one predictor using all three data types simultaneously (*f*_all_). To ensure a fair comparison, we split the training data of 10 461 MSAs into a *training set* and a *test set*. The *test set* is only used for performance comparison. Note that we trained a new predictor for this experiment instead of using `Pythia 2.0` to ensure that the predictor was trained on the *training set* only. The *training set* comprises 8368 MSAs and the *test set* contains the remaining 2093 MSAs. We split the data using stratified sampling such that the respective proportion of DNA, AA, and morphological MSAs is preserved. Thus *f*_DNA_ is trained on 7287 DNA MSAs, *f*_AA_ on 766 AA MSAs, and *f*_M_ on 315 morphological MSAs. *f*_all_ is trained on the full *training set* of 8368 MSAs (DNA, AA, and morphological). We then compare the MAE of the per-data-type prediction models to the prediction performance of *f*_all_ using the MSAs of the respective data type in the *test set*. For DNA and AA MSAs, the prediction accuracy is identical when only using DNA/AA MSAs for training, compared to using all three data types (MAE = 0.1). For morphological data, we observe a slight decrease in MAE for *f*_M_ (MAE = 0.12) compared to *f*_all_ (MAE = 0.11). We suspect that this is caused by the lack of sufficient training data. The predictor trained on only morphological MSAs was trained on only 315 MSAs making it more challenging for the predictor to generalize to unseen data. In contrast, the predictor using MSAs of all data types can detect patterns that are independent of the underlying data type of the MSA, and can thus learn more general difficulty characteristics using a substantially larger MSA collection.

#### 3.2.2 Feature Importance and Model Insights

Table 1 shows the feature importance for `Pythia 0.0` and `Pythia 2.0` respectively. Note that we computed the gain-based feature importance that directly measures the contribution of each feature to the performance improvement during model training. `Pythia 0.0` relied heavily on the maximum parsimony (MP) based features *RF*_MP_ and 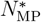 with a combined feature importance of approximately 75%. While `Pythia 2.0` still heavily relies on the *RF*_MP_ feature (31.5%), the most important feature is the Patterns-over-taxa ratio, with approximately 40%. We suspect that `Pythia 2.0` relies less on the 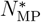 feature (5%), since the new training data shows an abundance of MSAs where 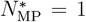, indicating that all 24 inferred MP trees have distinct tree topologies (see Figure 2).

**Table 1:**
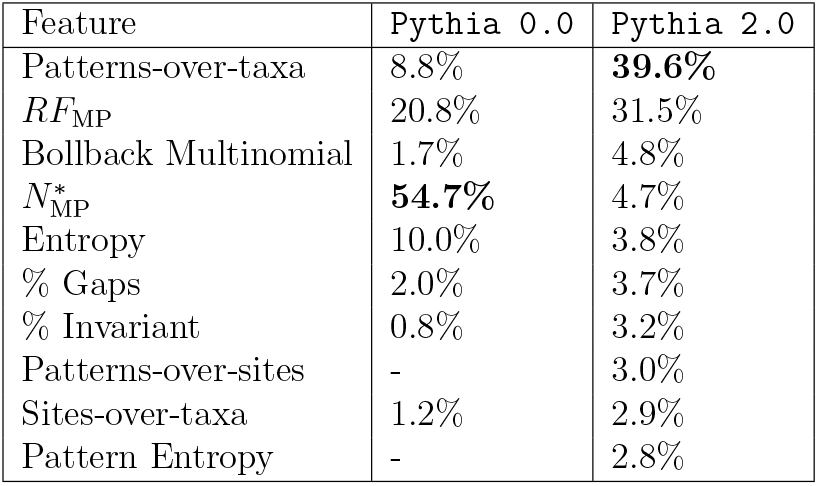
Feature importances in percent for Pythia 0.0 and Pythia 2.0. We computed the feature importances using the gain-based feature importance.

| Feature | Pythia 0.0 | Pythia 2.0 |
| --- | --- | --- |
| Patterns-over-taxa | 8.8% | **39.6%** |
| $RF_{MP}$ | 20.8% | 31.5% |
| Bollback Multinomial | 1.7% | 4.8% |
| $N_{MP}^*$ | **54.7%** | 4.7% |
| Entropy | 10.0% | 3.8% |
| % Gaps | 2.0% | 3.7% |
| % Invariant | 0.8% | 3.2% |
| Patterns-over-sites | - | 3.0% |
| Sites-over-taxa | 1.2% | 2.9% |
| Pattern Entropy | - | 2.8% |

Using *Shapley values* [15], we can gain additional insights into the contribution of individual features to the predicted difficulty. Shapley values are a method from cooperative game theory to fairly distribute the contribution of individual players to a total payout. In our case, the payout is the predicted difficulty for an MSA and the players correspond to the respective feature values. In contrast to feature importance, Shapley values consider cooperation effects between features based on all possible combinations of feature values. A feature’s contribution is determined based on the average change in predicted difficulty across all these coalitions. Shapley values are computed for a single prediction (the features of a single MSA and the corresponding predicted difficulty) and the interpretation of the resulting Shapley values only applies for the specific combination of feature values. For further details, please refer to the very intuitive and thorough explanation of Shapley values by Molnar [13].

`Pythia 2.0` includes a command line option, as well as a dedicated method to plot the Shapley values for a given MSA to allow for interpretable predictions, and supports Pythia users to gain additional insights. For these plots, as well as for the following analyses, we rely on the *SHAP* Python package [11].

Using the Shapley values computed on multiple MSAs, we can visualize a feature value contribution trend for the overall prediction via a so-called *beeswarm* plot. A beeswarm plot simply combines multiple, individual plots of Shapley values into a single figure. Figure 5 shows the beeswarm plot of `Pythia 2.0` for all 10 461 MSAs in our training data. The features are sorted by importance, with the topmost feature being the most important one. Note that the feature order differs compared to the feature importances stated in Table 1, since the feature importance is computed differently. The color of each dot corresponds to the magnitude of the respective feature value, with darker colors corresponding to higher values. The x-axis reflects the contribution of a feature value to the model output (the predicted difficulty). Negative values indicate a reduction in predicted difficulty, while positive values indicate a difficulty increase. The x-axis is relative to the *baseline* prediction. In the case of a GBT regressor, this baseline prediction simply corresponds to the average difficulty in the training data.

**Figure 5:**
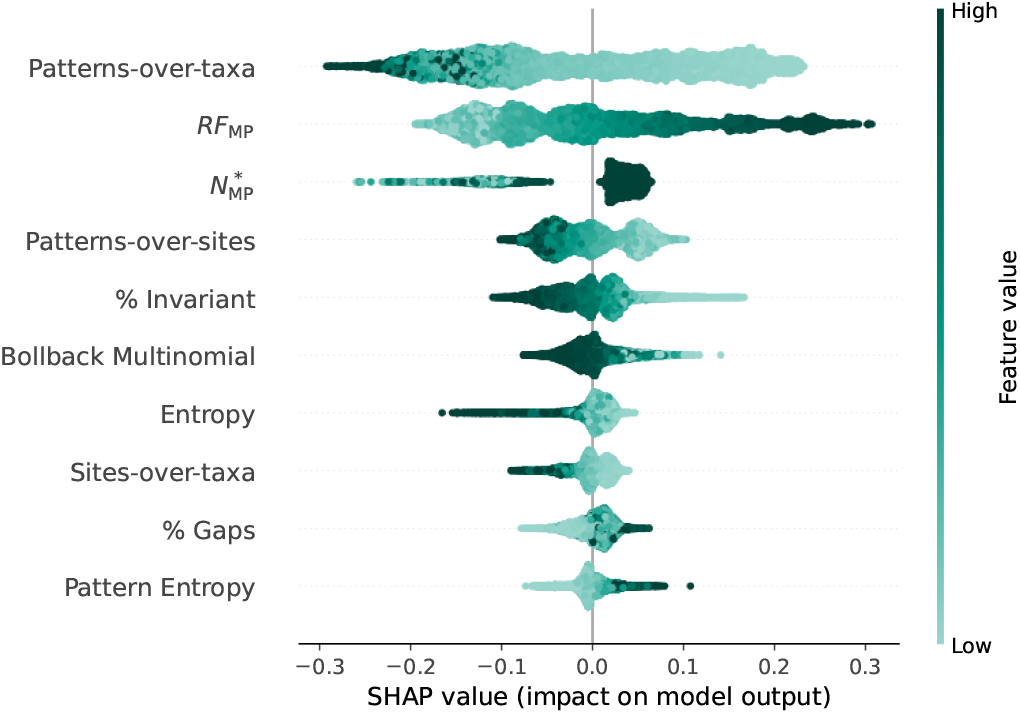
Beeswarm plot of Shapley values of `Pythia 2.0` for the 10 461 MSAs in our training data.

The beeswarm plot clearly shows that a higher signal in the MSA corresponds to lower predicted difficulties. This is predominantly visible for the patterns-over-taxa ratio, with contributions as low as −0.3. This trend is also visible for the patterns-over-sites ratio, the sites-over-taxa ratio, and the proportion of gaps. Analogously, a higher information content (as captured by the Entropy, the Bollback Multinomial, and the Pattern Entropy) has a negative impact on the prediction, thus contributing towards an MSA being easier to analyze. Note that, while *higher* values indicate a *higher* information content for the Entropy and the Bollback Multinomial, *lower* values indicate a *higher* information content for the Pattern Entropy. Interestingly, a *higher* proportion of invariant sites contributes *negatively* to the predicted difficulty and vice versa. This is unexpected, since a high proportion of invariant sites is expected to induce less variation (and thus signal) for phylogenetic inference. Consequently, we would expect higher proportions of invariant sites to induce higher predicted difficulties. Further investigating this counter-intuitive trend will be the subject of future work. As expected, higher *RF*_MP_ values contribute to an MSA being predicted as more difficult, while lower values contribute to predicting lower difficulties. This trend is expected, as the RF-Distance between the MP trees is a good indicator for the RF-Distance between ML trees, which is directly related to our quantification of the ground-truth difficulty (see Haag *et al*. [5]). Analogously, a higher proportion of unique topologies 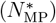 tends to contribute positively to the predicted difficulty. However, this trend is not as clearly pronounced. We again suspect that this is caused by the high abundance of MSAs where 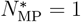.

#### 3.2.3 Runtime Comparison

In the following, we benchmark the runtime of `Pythia 2.0` against `Pythia 0.0`, and against ML tree inference using RAxML-NG. Additionally, we provide an overview of the run time contributions of MSA parsing, feature computation, and difficulty prediction to the overall difficulty prediction runtime for a single MSA in `Pythia 2.0`. We used all 10 461 MSAs in our training dataset for all subsequent analyses.

`Pythia 2.0` is on average 2.4 ± 1.2× faster than `Pythia 0.0` (2.0×median). Multiple factors contribute to this speedup. Firstly, we reduced the number of MP trees in `Pythia 2.0` to 24, compared to the 100 MP trees in `Pythia 0.0`. We also refactored the internal representation of MSAs in `Pythia 2.0`. This allows us to natively compute the number of patterns, the proportion of invariant sites, and the proportion of gaps in Python using our MSA representation. In contrast, in `Pythia 0.0`, we relied on RAxML-NG for computing these features. While the RAxML-NG-based implementation is still faster, including these features as MSA properties in `Pythia 2.0` (and thus removing the RAxML-NG dependency) improves the re-usability of our MSA representation in our PyPythia Python library for other applications relying on MSA attributes. We observe that the speedup of `Pythia 2.0` compared to `Pythia 0.0` increases with increasing MSA size (number of taxa times number of patterns in the MSA; see Figure 6). For instance, the average speedup for MSAs with a size that exceeds 100 000 is 4.2× (4.1× median).

**Figure 6:**
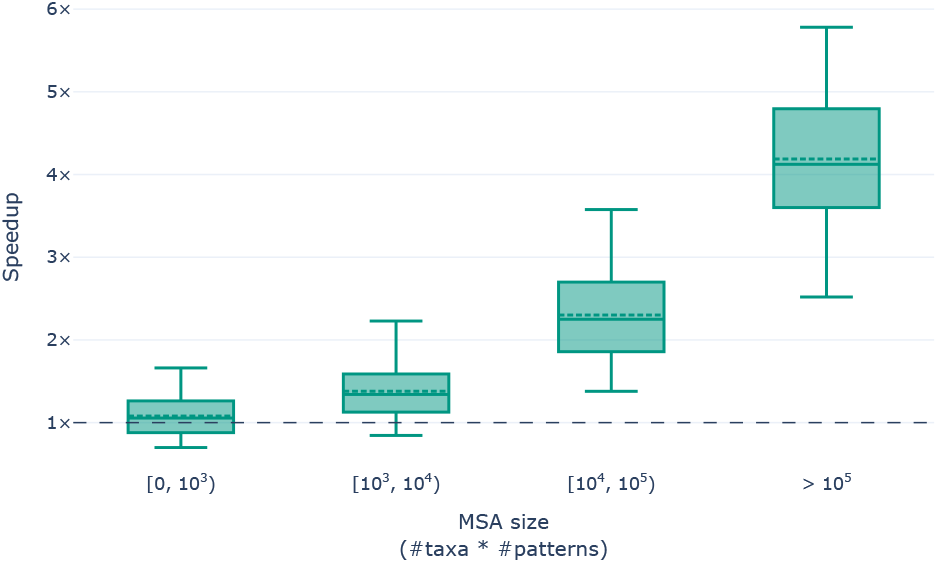
Speedup of `Pythia 2.0` compared to `Pythia 0.0` as a function of MSA size. MSA size is computed as the number of taxa times the number of unique site patterns in the MSA. To improve visualization, we only show data between the 5th and 95th percentile.

The main goal of using Pythia is to gain insights into the expected behavior of the MSA under ML phylogenetic inference. This allows for a more informed analysis setup and adjustment of expectations before conducting any time- and resource-intensive analyses. To demonstrate the value of Pythia, we thus compare the runtime of predicting the difficulty for an MSA using `Pythia 2.0` to the runtime of performing a *single* ML tree inference using RAxML-NG. Across all MSAs, we observe an average speedup of 21.5 ± 43.8× (7.7× median). We observe a high standard deviation since the speedup predominantly depends on the MSA size. Figure 7 shows the speedup as a function of MSA size. For (very) small MSAs with a size below 1000, `Pythia 2.0` is slightly slower than a single ML tree inference. However, the average speedup for MSAs with a size exceeding 100 000 is above 50× (37× median). Note that an MSA with 100 taxa and 1000 patterns already falls into this category.

**Figure 7:**
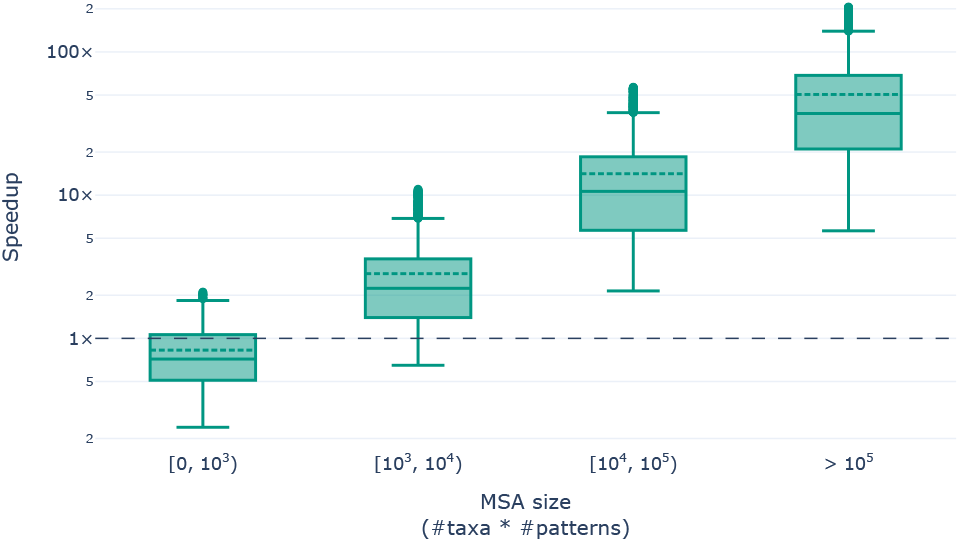
Speedup of `Pythia 2.0` compared to a single ML tree inference using RAxML-NG as a function of MSA size. The MSA size is computed as the number of taxa times the number of unique site patterns in the MSA. Note that the y-axis is logarithmic. To improve visualization, we only show data between the 5th and 95th percentile.

Finally, we analyzed the contribution of various computational steps required for difficulty prediction in `Pythia 2.0` with respect to the overall runtime. Figure 8 shows the average contribution across all MSAs in our training data. Note that we summarized the feature computation into three separate groups: the *MP-based* group (*RF*_MP_ and 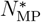), the *Information Content* group (Entropy, Bollback Multinomial, Pattern Entropy), and the *MSA Attributes* group (sites-over-taxa, patterns-over-taxa, patterns-over-sites ratio, % invariant, % gaps). *MSA Parsing* captures the runtime for reading the MSA from file and transforming it into our internal MSA representation. *Prediction* refers to loading the pretrained LightGBM model and performing the final difficulty prediction. The runtime is dominated by the computation of the MP-based features, with an average contribution of 67%. This indicates that future work in Pythia should focus on improving the runtime of the MP tree inference, either by improving the MP implementation in RAxML-NG (which is developed in our research group) or by replacing RAxML-NG with a faster alternative.

**Figure 8:**
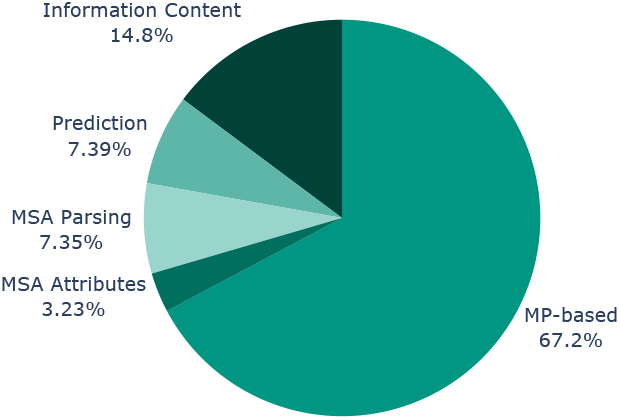
Contribution of computational steps in `Pythia 2.0` to predict the difficulty of an MSA. The pie chart shows the average contribution across the 10 461 MSAs in our training dataset.

## 4 Ground-Truth Labeling Tool

Computing the ground-truth difficulty for an MSA is time- and resource-intensive. As presented in Haag *et al*. [5], we quantify this difficulty based on 100 ML tree inferences using RAxML-NG [10], and a subsequent filtering of trees using the statistical tests implemented in IQ-TREE [12]. To train Pythia, we implemented a custom pipeline using the Snake-make workflow management system [9] to generate the ground-truth labels for many MSAs in parallel (see Haag *et al*. [5] for details). However, setting up and running this pipeline is somewhat complicated, as it is intended to generate the labels for many MSAs in one run using our institutional cluster. To simplify the computation of the ground-truth difficulty for a single MSA, we implemented a new command-line tool called *Pythia Difficulty Label Generator* (*PyDLG*). PyDLG and its corresponding documentation are available as open-source code via GitHub at https://github.com/tschuelia/PythiaLabelGenerator. PyDLG takes an MSA as input and performs all required tree inferences, statistical tests, and RF-Distance computations to obtain the ground-truth difficulty. While our original quantification is based on 100 ML trees, we additionally allow the user to adjust the number of inferred trees. Please note that using less than 100 trees will only approximate the true difficulty.

As we have shown above, Pythia is on average approximately 20 times fast than a *single* RAxML-NG tree inference. We thus strongly encourage users to predict the difficulty of their MSA using our highly accurate `Pythia 2.0` difficulty predictor instead of PyDLG. `Pythia 2.0` is trained on DNA, AA, and morphological data, and we are confident that Pythia can accurately predict the difficulty for any biological MSA. Consequently, we suggest users to only use PyDLG if they are convinced that Pythia is unable to correctly predict the difficulty of their MSA. We do note that this appears to be the case for MSAs that are used to infer phylogenies of natural languages (currently unpublished work by our research group member Luise Häuser).

## Conclusion

Our latest difficulty prediction model `Pythia 2.0` was trained on approximately three times more MSAs than our initial version. Since we include morphological MSAs in addition to DNA and AA data, `Pythia 2.0` can now accurately predict the difficulty for all three data types. We introduced two new features, reduced the number of required MP trees, and trained a new machine learning model. These changes result in a slight prediction accuracy improvement. For predicting the difficulty on a scale from 0 to 1, `Pythia 2.0` is highly accurate with a MAE of 0.08 (MAPE = 2.2%). Compared to our initial version `Pythia 0.0`, `Pythia 2.0` is on average 2.4 times faster. We showed that the MP-based feature calculations dominate the overall runtime of `Pythia 2.0`. Thus, future work should focus on runtime improvements of the MP tree inference. With PyDLG, we additionally presented a new and easy-to-use command line tool to compute the ground-truth difficulty for a given MSA.

## Acknowledgements

This work was financially supported by the Klaus Tschira Foundation, by the European Union (EU) under Grant Agreement No 101087081 (Comp-Biodiv-GR).

---

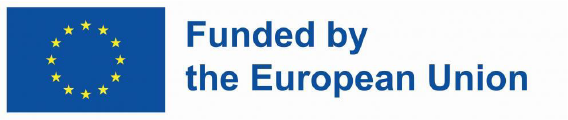

## References

[1] T. Akiba, S. Sano, T. Yanase, T. Ohta and M. Koyama. Optuna: a next-generation hyperparameter optimization framework. in Proceedings of the 25th ACM SIGKDD International Conference on Knowledge Discovery & Data Mining, pages 2623–2631, New York, NY, USA, 2019. ACM.

[2] L. Breiman. Random forests. Machine Learning, 45(1):5–32, 2001.

[3] L. R. Foulds and R. L. Graham. The steiner problem in phylogeny is NP-complete. Advances in Applied Mathematics, 3(1):43–49, 1982.

[4] J. H. Friedman. Greedy function approximation: a gradient boosting machine. The Annals of Statistics, 29(5), 2001.

[5] J. Haag, D. Höhler, B. Bettisworth and A. Stamatakis. From easy to hopeless – predicting the difficulty of phylogenetic analyses. Molecular Biology and Evolution, 39(12), 2022.

[6] D. Hoehler, J. Haag, A. M. Kozlov and A. Stamatakis. A representative performance assessment of maximum likelihood based phylogenetic inference tools. bioRxiv, 2022.

[7] D. Höhler, W. Pfeiffer, V. Ioannidis, H. Stockinger and A. Stamatakis. RAxML Grove: an empirical phylogenetic tree database. Bioinformatics, 38(6):1741–1742, 2021.

[8] G. Ke, Q. Meng, T. Finley, T. Wang, W. Chen, W. Ma, Q. Ye and T.-Y. Liu. LightGBM: a highly efficient gradient boosting decision tree. in Proceedings of the 31st International Conference on Neural Information Processing Systems, NIPS’17, pages 3149–3157, Red Hook, NY, USA, 2017. Curran Associates Inc.

[9] J. Köster and S. Rahmann. Snakemake – a scalable bioinformatics workflow engine. Bioinformatics, 28(19):2520–2522, 2012.

[10] A. M. Kozlov, D. Darriba, T. Flouri, B. Morel and A. Stamatakis. RAxML-NG: a fast, scalable and user-friendly tool for maximum likelihood phylogenetic inference. Bioinformatics, 35(21):4453–4455, 2019.

[11] S. M. Lundberg and S.-I. Lee. A unified approach to interpreting model predictions. In I. Guyon, U. V. Luxburg, S. Bengio, H. Wallach, R. Fergus, S. Vishwanathan and R. Garnett, editors, Advances in Neural Information Processing Systems 30, pages 4765–4774. Curran Associates, Inc., 2017.

[12] B. Q. Minh, H. A. Schmidt, O. Chernomor, D. Schrempf, M. D. Woodhams, A. von Haeseler and R. Lanfear. IQ-TREE 2: new models and efficient methods for phylogenetic inference in the genomic era. Molecular Biology and Evolution, 37(5):1530–1534, 2020.

[13] C. Molnar. Interpretable Machine Learning. 3 edition, 2025.

[14] W. H. Piel, L. Chan, M. J. Dominus, J. Ruan, R. A. Vos and V. Tannen. TreeBASE v.2: a database of phylogenetic knowledge. e-BioSphere 2009, 2009.

[15] L. S. Shapley. 17. A value for n-person games. in Contributions to the Theory of Games (AM-28), Volume II, pages 307–318. Princeton University Press, 1953.

[16] A. Togkousidis, O. M. Kozlov, J. Haag, D. Höhler and A. Stamatakis. Adaptive RAxML-NG: accelerating phylogenetic inference under maximum likelihood using dataset difficulty. Molecular Biology and Evolution, 40(10), 2023.

